# Non-destructive monitoring of tissue-specific, irrigation responsive impedance signals in leaves using microneedles probes

**DOI:** 10.1101/2025.03.31.646420

**Authors:** Kaitlyn J.H. Read, Patrick J. Hudson, Jose Monclova, Haley Monteith, Ronen Polsky, David T. Hanson, Philip R. Miller, Kati Seitz

## Abstract

A non-destructive methodology for monitoring impedance changes in sorghum leaves was developed and recorded irrigation-dependent responses that differed between leaf tissues. Metal microneedles were used as impedance probes and were shown to cause minimal damage to the plant. The needles were placed on either the abaxial or adaxial side of the leaf midrib using small clamps and re-used hundreds of times with minimal signs of wear. Cross-sectional images verified the precision of microneedle placement near vascular bundles on the abaxial surface and in non-vascular hydrenchyma on the adaxial surface. Impedance measurements with microneedles displayed a significant decrease in resistance compared to planar electrodes due to bypassing the epidermal layer. A tissue-specific impedance response was seen in relation to irrigation where the non-vascular adaxial surface remained largely stable throughout a day of measurement, while impedance increased in the vascular abaxial surface during exposure to light and decreased following watering. Impedance data were also compared with simultaneous gas exchange measurements of photosynthesis and transpiration.

## Introduction

Precision agriculture is enabling a new generation of crop management by supplementing traditional methods of growing with spatial and temporal dynamics data supplied by crop and weather monitoring technology, resulting in greater yield and lower irrigation (Sishodia et al., 2020). One area of great interest in this field is non-destructive methods for real-time monitoring of plant water potential (the potential energy of water per unit volume used to determine if water will move from one area to another). However, development remains a challenge, resulting in a potential invaluable stream of data regarding plant health and stress undermined and undervalued (Levinsh, 2023). Standard methods for measuring plant water status (*i.e*., water potential) involve destructive sample collection and are not amenable to real-time monitoring (Boyer, 1967; Boyer, 1966; Greenwood et al., 2010). Soil-based sensors, a non-destructive alternative to monitoring plant water status, offer insight into localized soil water content and are used in feedback loops of commercially available irrigation management systems, which has resulted in water savings at crop scales (van Iersel et al., 2009; Koman et al., 2017). While these sensors are simple, inexpensive, and reusable, their placement both in depth and proximity to the plant can affect the ability to accurately predict plant water requirements. Ultimately, on-plant measurements could offer better insight into plant health and water status, however, these technologies have been challenging to implement. Wearable sensors are gaining popularity due to their ability to provide valuable health information in a non-invasive manner and various efforts are emerging for agricultural exploration of these devices (Yin et al., 2021). The commercial market for precision agriculture sensors has expanded to include many wearable and wireless sensors (Shafi et al., 2019, Reynolds et al. 2023). These devices have provided unique insight into plant functions previously either challenging to measure or not possible to continuously measure in the field.

Many wearable sensors utilize new materials or advanced fabrications methods, but few have attempted to apply these advances to basic studies of plant physiology using electrochemical impedance spectroscopy (EIS). EIS is an AC electrochemical measurement that monitors resistance and phase angle changes of a material (biotic or abiotic) across a range of frequencies (∼0.1-100kHz) and uses a representative electrical circuit of the material to model fundamental properties. When EIS is used to study biological systems, correlations of electrical components in the representative circuit (i.e. resistors, capacitors) are made with their biological component analogs (i.e. extracellular fluid, cell membrane) to draw conclusions on how changing environmental conditions affect these biological components (Azzarello et al., 2012). In principle, the electronics to run an EIS measurement are simple and inexpensive, and their low current operation makes them safe for medical usage. Correlations between EIS data and plant health states have been demonstrated in heat and cold stress (Repo et al., 2000), fruit ripening (Varlan et al., 1996), biomass (Bragós et al., 1999), and water status (Jamaludin et al., 2015). In addition, the hardware to operate these systems has shrunk such that field-deployable units are both feasible and economical (Ferreira et al., 2010).

Probes (signal transducers) for plant EIS measurements come in a range of formats and are largely dependent on their application. Metal rods in soils for measuring plant root biomass, metal clips for anchoring to plant stems, and wires/needles for in situ measurements have all been extensively demonstrated (Repo et al., 2005; Jeon et al., 2017). We look to adapt microneedle technology as an EIS probe for plant leaf measurements due to their defined size, strength, and demonstrated minimally invasive nature. Given that leaves are the primary plant interface for gas exchange, measurement of this tissue may be advantageous from a sensing perspective, however, they are also a particularly challenging sample due to their delicateness compared to the rest of the plant. Microneedle technology, which was initially developed for drug delivery and has been adapted for wearable sensing applications, uses short (<1 mm) high-aspect ratio needles to penetrate the skin’s protective barrier, and is considered a non-invasive technology due to the depth of puncture (Kim et al., 2012; Miller et al., 2016). Several examples exist of coupling microneedle technology with EIS in humans (Li et al., 2016) and more recently in plant stalks (Jeon et al., 2017; Garlando et al., 2022). In this study, we investigated if metal microneedles EIS probes could be selectively placed in different leaf anatomy of *S. bicolor* and if these specific tissues exhibited different water dependent responses given our understanding of their physiological function.

## Methods

### Microneedle fabrication

Microneedle fabrication was performed using a previously developed method where hollow metal microneedles were created via a laser direct write (LDW), Polydimethylsiloxane (PDMS) micromolding, and electroplating (Miller et al., 2019). Hollow needles were not needed for this application; however, this microneedle fabrication methodology was chosen due to strength and conductivity of the created materials. In brief, two-photon lithography with LDW was used to create a solid microneedle based on a CAD file that served as the master structure for subsequent PDMS molding. The microneedle was a four-sided pyramid that stood 700 μm tall and 220 μm wide at the base. A Ti-Sapphire laser operating at 780 nm and 150fs was used as the light source to polymerize a photosensitive resin (Eshell300), which was positioned over a substrate made of the same material such that polymerization would adhere the two components.

Location and pattern of polymerization was controlled by software (GOLD3D), which converted the CAD files into individual layers for LDW, and a three-axis stage. Following the LDW process, parts were washed with isopropyl alcohol (IPA) and exposed to UV light for 15 min to complete the curing process. Molds of the master structures were made by casting PDMS (Sylgard 187) at varying ratios of catalyst to precursor over the solid microneedles and then baked at 70C for 1 hour. Molds were removed and coated with a metal seed layer (100A titanium/1000A gold) using electron beam physical vapor deposition. Metal coated molds were then electroplated in either a nickel or iron bath to backfill the mold. Following electroplating, the electroformed microneedles were removed from the molds by hand.

### Plant material

*Sorghum bicolor* (Silage Master) were grown in an 4>×8 Grow Tent S848 (Vivosun Inc., Ontario, California). Seeds were germinated in 2.6 L pots and watered with 150 mL of water daily, then transferred to 8.3 L pots filled with Mastermix-820 (Sun Gro Horticulture, Seba Beach, AB, Canada) after 6 weeks and received 500 mL of water daily. Plants experienced LED artificial daylight from 6:00 am to 8:00 pm; light intensity ranged from 800 μmol m^-2^ s^-1^ at the base of the lights to 300 μm m^-2^ s^-1^ at the level of the soil. The growth tent temperature was held at 25 °C during light hours and 21 °C during dark hours. Plants were measured when they had 9 fully expanded leaves but before flowering. The mid-rib was chosen as the location for microneedle insertion due the clear distinction of tissue types (i.e. vascular and hydrenchyma) that are critical to plant function, the ease of access to each tissue, and the spatial differentiation of the tissues.

### Microneedle insertion into tissues

A microneedle array was composed of 2 microneedle probes with 2 needles fabricated 500 μm apart to prevent pivoting on the plant. The two probes were spaced 6 cm apart and were positioned on the midrib of either the adaxial or abaxial surface of the leaf. Two microneedle probes were pressed on the adaxial surface of the midrib until the microneedles punctured the epidermal tissue to measure the hydrenchyma. The probes were held in place with a clamp or tape. This process was repeated on the abaxial surface to measure tissues around the vascular bundles.

To verify placement and assess wounding, microneedle probes were coated in safranin red, then inserted into the adaxial and abaxial surfaces of midribs. Microneedles were immediately removed to prevent staining of the tissues surrounding the immediate inject site. After the probe was removed, midribs were cross-sectioned, and the puncture sites were imaged using a CCD camera and dissecting microscope (Zeiss Axioscope, Carl Zeiss, Germany). We visually assessed adverse effects of microneedle placement over time on the plant and the effects on microneedle morphology in response to repeated insertions into plant tissues. Effects on plants were examined by placing arrays in both midrib tissues on a fully expanded leaf, secured by wooden clothespins. After 14 days, the puncture site was identified and imaged. Repeated insertion effects on microneedle morphology caused by puncturing the cell walls of the midrib were tested with an unused microneedle probe. The probe was inserted into the midrib 1, 5, 10, 50, and 100 times (cumulative) and imaged under a dissecting microscope to inspect for damage to the microneedle structure.

### Electrical Impedance Spectroscopy

1 x 2 Microneedle arrays were used for each tissue-specific measurement with two probes placed 6 cm apart in the midrib on the abaxial surface for measurements in the hydrenchyma, or on the abaxial surface for vascular measurements (Supplemental Figure 1). Microneedles were placed along the midrib of the third fully expanded leaf from the top of the primary shoot. The first probe was placed near the shoot, distal to the ligule. The second probe was placed 6 cm distal to the first probe along the midrib. To demonstrate the necessity of subdermal access for EIS measurements, planar foil lacking microneedles was attached to the surface of the midrib without puncturing the epidermal layer. Planar foil probe arrays were placed 6 cm apart along the abaxial and adaxial surface of the midrib, distal to the microneedle arrays. The planar foil probes were manufactured under the same conditions with the same materials as the microneedle arrays.

Using a handheld LCR meter, point measurements of impedance (Z, kΩ) and phase-shift (Θ, °) in the hydrenchyma, vascular tissues, and epidermis were made at three different frequencies (0.1, 1, and 10 kHz). Impedance measurements were collected with a LCR meter (BK Precision) by passing a 0.6 V AC between an array set. Data was collected hourly from 9:00 to 13:00. Plants were watered with 250mL of water immediately following 13:00 data collection. Data was then collected every 30 min from 13:30 – 15:00 pm. Three replicates were recorded from the LCR meter. The mean of the three replicates for each frequency was calculated as the value for each time point.

### Impedance Calculation

Values of impedance from the LCR meter were taken in impedance magnitude and phase angle. Determining real and imaginary components of impedance and the capacitance were determined using the following equations where Z is the magnitude of impedance; angle is the phase shift between the voltage and the current; R is resistance of the AC circuit (Real component); I is the reactance of the AC circuit (Imaginary component); C is the capacitance of the dielectric; j is the square root of –1; and w is the angular frequency.

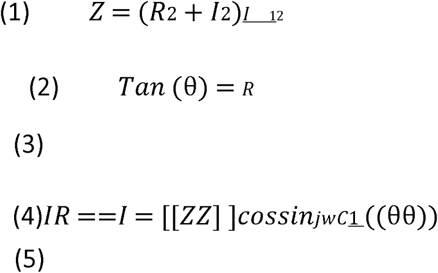

The real and imaginary components of impedance are the Cartesian coordinates of the complex number.

### Plant gas exchange

Net assimilation (A_net_,μmol m^-2^ s^-1^), transpiration (E, mol m^-2^ s^-1^), and stomatal conductance (g_sw_, mol m^-2^ s^-1^) were measured with a LiCor 6800 infrared gas analyzer, which was attached distal to microneedle arrays on the same leaf. Gas exchange measurements were performed in tandem with the microneedle impedance measurements to determine if additional insight into the insertion site specific response could be gathered from additional characterization. Chamber conditions were set to reference [CO_2_] to 400 ppm, vapor pressure deficit to 1.8 kPa, temperature to 25 °C, and photosynthetically active radiation (PAR) to 1500 μmol m^-2^ s^-1^. Leaf tissue in the LiCor 6800 chamber was allowed to stabilize for 30 minutes before measurements were collected. Data were collected concurrently with EIS data.

### Data analysis

All analysis and graphs were compiled based on the averaged impedance value calculated in R and the single value gas exchange data from the LiCor 6800. Standard error was calculated in R. All graphs were generated in R using the following packages: ggplot2, scales, ggplotgui, viridis, lubridate, lemon, plyr, and gridExtra. Statistically significant changes in response variables through time were identified using a linear mixed effects modeling approach (LMM). Model construction and analyses were carried out using R packages: lme4, lsmeans, lmerTest, multcomp, pbkrtest.

## Results

### Microneedle Positioning

In this study, a microneedle-based plant leaf EIS sensor was investigated for measurements of specific plant leaf tissue responses under varying water conditions. The leaf anatomy within the midrib of a *S. bicolor* is vertically differentiated as seen in the cross-sectional images of Supplementary Figure S2 and provides a unique testbed for investigating our tissue-specific hypothesis. Two main structural tissues are present, hydrenchyma and vascular bundles, within the S. *bicolor* leaf midrib. Hydrenchyma is a water storage tissue primarily located in the upper section of the midrib, but also extends to the abaxial surface, that provides mechanical support and water storage within the leaf. Vascular bundles contain the xylem and phloem, which transports water from the roots to the leaves and sugars from the leaves to the roots respectively. Because the primary functional xylem water transport cells are dead, void of cell contents at maturity, and have large perforations through the walls at cell ends, it is able to move water from the roots to the leaves with very little resistance.

Our first experiments using microneedles on plants explored placement and attachment of microneedles on plant leaves and how the plant responded. Positioning the microneedles on the mid-rib was easily accomplished by hand in a repeatable manner without assistance from optical magnification or micro-manipulators. Cross-sectional slices of microneedle punctured midribs were performed with dye puncture (Figure 1 A & B) via dip coating microneedle arrays prior to insertion. This technique was utilized to understand depth of microneedle puncture, which can be elusive in more elastic tissue, and for identifying puncture locations when performing cryo-cross sections. While elucidating depth of puncture in these experiments was challenging, the images highlight that the area surrounding the punctured tissue remains largely unchanged, which may be indicative of the non-invasive nature of the probes. Optical images following hand application confirmed insertion location (Figure 1 C & D) via the indentation left after microneedle puncture and indicated that point-based EIS measurements could be performed quickly and accurately within the tissue of interest. For continuous EIS measurements, wooden laundry clips were used to hold the microneedle transducers in position. The combined weight of the microneedle array and the laundry clip didn’t cause adverse effects. The microneedles were removed after 14 days and the insertion site was identified and examined for adverse response (Supplementary Figure S3). Minimal damage was seen surrounding the insertion site and punctures were identified by slight discoloration compared to the surrounding tissue, which may be due to localized cell death at the epidermis. We suspect that the pressure from the clamps may lead to the localized discoloration since measurements performed with clips with less compressive force didn’t produce such a response. Previous studies using wires in leaves indicated a small collar of damage at the insertion site of plant tissue that didn’t disrupt local or system level function (Zhang and Wilson, 1993). These initial results indicate that adhering microneedles to plants is well tolerated over 14 days and that placement of metal microneedles within leaf tissues does not initiate a severe adverse response.

**Figure 1:**
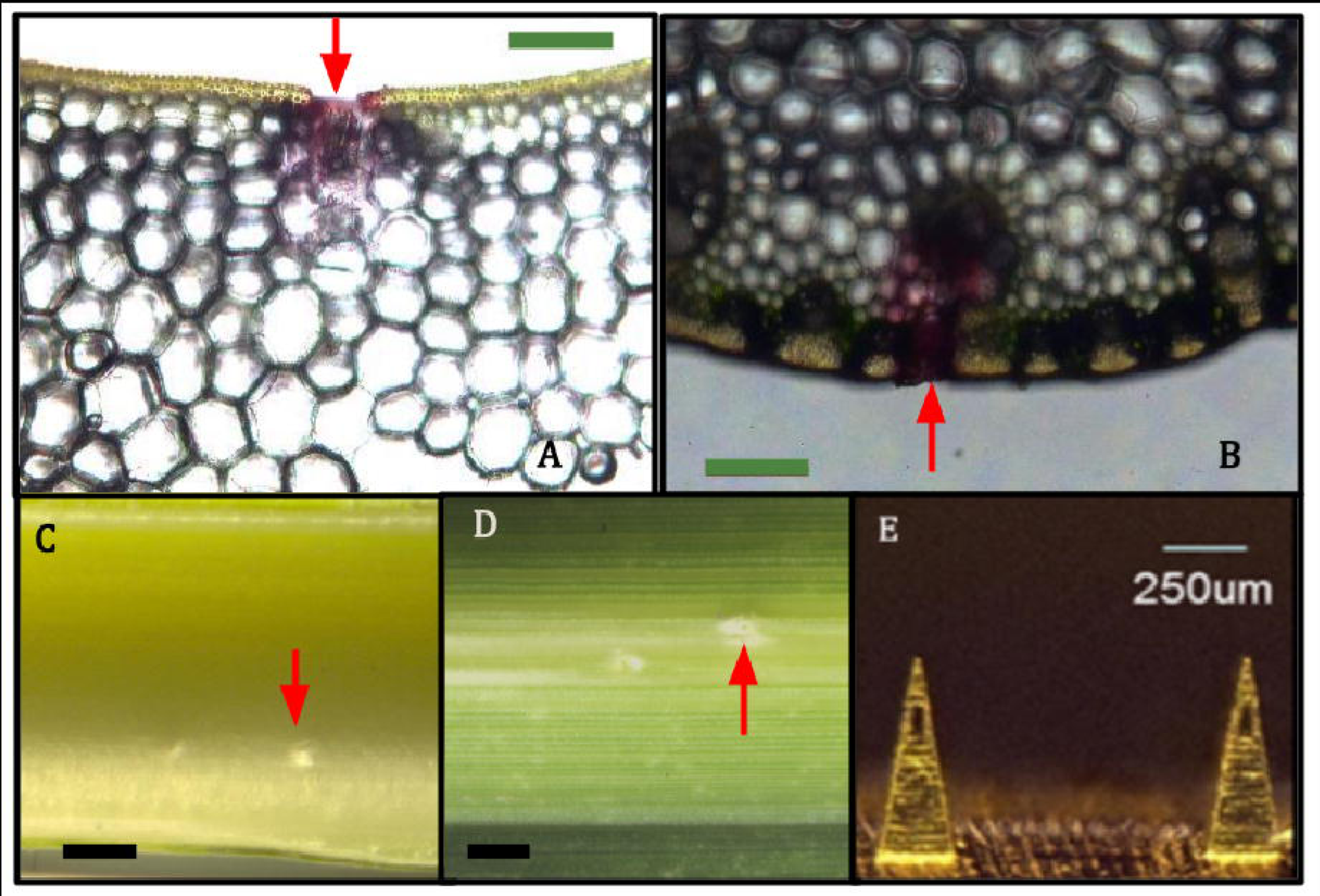
Microneedle insertion sites. **(A)** Optical image of the adaxial surface of a *S. bicolor* sectioned perpendicular to the midrib showing safranin red stain left by a microneedle insertion in the hydrenchyma; **(B)** Optical image of the abaxial surface of a *S. bicolor* sectioned perpendicular to the midrib showing safranin red stain left by a microneedle insertion in the vascular bundle; **(C)** Optical image of the adaxial epidermal surface of a punctured *S. bicolor* midrib with no stain; **(D)** Optical image of the abaxial epidermal surface of a punctured *S. bicolor* midrib with no stain; **(E)** Optical image of a representative microneedle array showing the two examples of similar microneedles. Although not the needles used for the study, the manufacturing techniques and geometeries are similar to the ones used. Scale bars: 500 µm (A, B, C, D); 250 µm (E). These are representative images of 5 replicates.

### Microneedle Puncture Kinetics and Durability

Analysis of microneedle insertion is typically performed by taking cross-sectional images of insertion sites to determine puncture kinetics, possible needle fracture, and compaction of surrounding tissue during insertion, which can limit performance of microneedle use (Wang et al., 2006). To our knowledge, microneedles insertion kinetics into plant tissue has not been studied, so cross-sectional images were taken (Figure 1 A & B) following insertion into both the adaxial and abaxial surface of sorghum midribs. In both insertion sites, the epidermal layer is not present, and the width of the void is comparable to the width of the microneedle base. Tissue compaction or tenting of the surrounding tissue was not detected; in animal skin, these issues can limit microneedle puncture depth and the desired placement. However, due to the rigidity of the midrib and the width of the insertion site, those issues are not seen in this application, and thus we can assume that nearly the entire length of the needle penetrated the plant tissue. The microneedles are large enough to visual detect needle fracture, and that was not observed in this study. These results indicate that microneedles can be reliably placed into desired tissue compartments, with no unintended breaching into adjacent tissues. (Figure 1 A & B). For point-based measurements, we explored the robustness of the metal microneedles by performing multiple insertions while documenting their structure integrity and shape. After 100 punctures (Figure 2) minimal deformation was seen. Subsequent testing revealed no noticeable impact on ease of insertion, microneedle integrity, or signal integrity following upwards of 700 punctures.

**Figure 2:**
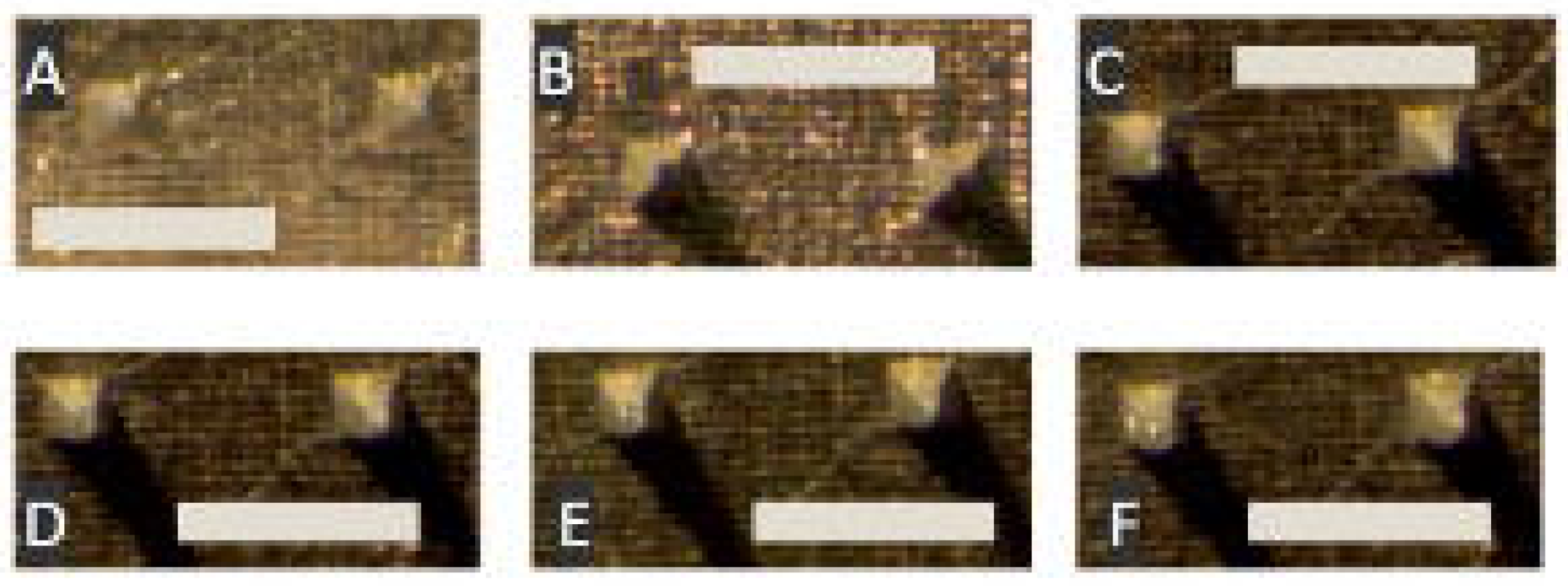
Microneedle arrays after puncturing a S. bicolor midrib. **(A)** A microneedle array before puncturing a midrib; **(B)** after a single puncture; **(C)** after five punctures; **(D)** after ten punctures; **(E)** after fifty punctures; and **(F)** after one hundred punctures. These are representative images of 5 replicated series of tests. 15X Magnification for all images, 1mm scale bar.

### Tissue-specific impedance

In our EIS application, we theorized that by bypassing the waxy cuticle and epidermis of plant tissue, the background resistance of the impedance measurement that did not contribute to the underlying physiology would be decreased or removed, enabling high-resolution measurements of the tissue of interest. To test this theory, planar metal electrodes were tested in conjunction with microneedle EIS probes, and impedance measurements were performed using our hand-held impedance analyzer at frequencies of 0.1, 1, & 10kHz (Figure 3). Probe placement for these tests, and for the continuous tests, had both probes (i.e. signal out and signal in) placed on the same side of the leaf (abaxial or adaxial) 6 cm apart. Surface area and material (Gold – Au) of the planar electrodes matched that of the microneedle arrays (Figure 2) so we assumed the only difference in measurement was the microneedles. When compared to the impedance values taken with microneedle EIS probes, impedance magnitude of the planar electrodes were 2 orders of magnitude higher at low frequencies (0.1 and 1.0 kHz, Figure 3 A and C) and one order of magnitude higher at our high frequency (10 kHz, Figure 6 D) test for both abaxial and adaxial measurements. The planar electrode configuration had significantly more error and may be attributed to the lack of conductive gel that typically is used to electrically couple to the test specimen.

**Figure 3:**
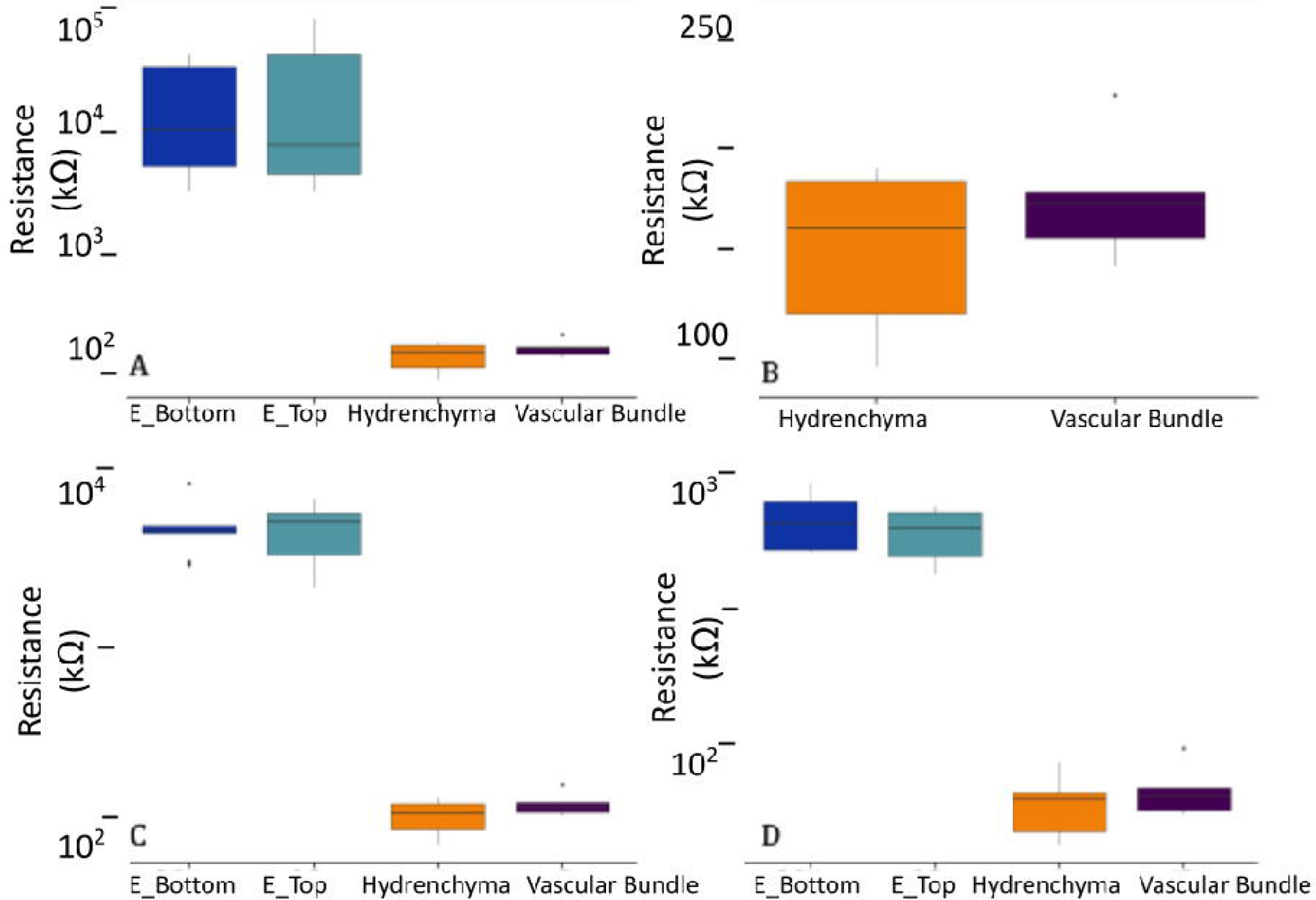
Magnitude of Impedance at the start of the experiment. The mean resistance of arrays *S. bicolor* leaf midribs. Box plot of the mean resistance of microneedle and planar arrays on the abaxial and adaxial surface of a *S. bicolor* midrib at 9am. **(A)** 0.1 kHz (Start of measurements) including epidermal measurements as shown on a log scale. **(B)** Box plot of microneedle on the abaxial and adaxial surface of a S. bicolor midrib at 9am at 0.1 kHz as shown on a linear scale; This is a closed look at the data from the microneedle probes. **(C)** 1 kHz as shown on a log scale; and **(D)** 10 kHz as shown on a log scale. Error bars represent 1 standard error, n=5, all measurements collected at 9 am.

The significant decrease in impedance gained by using microneedles as EIS probes over planar electrodes (Figure 3 A & B) justified our hypothesis regarding epidermal associated resistance and encouraged investigation of tissue-specific impedance responses. To test this theory of tissue specific response, we set up a time course measurement where S. *bicolor* would go through a period of solar irradiation in a greenhouse while withholding water followed by saturating the soil with water (Supplemental Figure S4, Figures 4 & 5). In this test setup, microneedles were attached to the leaf midrib with laundry clips throughout the course of measurement. Potted plants under investigation were well watered and allowed to come to equilibrium for one hour prior to starting the experiment. Impedance and phase angle measurements were performed every hour and at every half-hour following the 250 mL irrigation at hour 4. Initial impedance and phase angle values of each microneedle measurement exhibited similar values, while planar electrode measurements had significantly higher impedance magnitudes at all frequencies at 0hr (Figure 4). As the time course study progressed and soil moisture pools diminished and ambient temperature increased, obvious differences in impedance magnitudes between the adaxial and abaxial surface were detected at all frequencies (Figure 5A, B, C). After four hours, the abaxial measurement increased by ∼250% at 0.1kHz while the adaxial measurements remained nearly constant, with no significant differences between subsequent time points. Planar transducer measurements showed random fluctuations in impedance for each surface tested (Figure 4).

**Figure 4:**
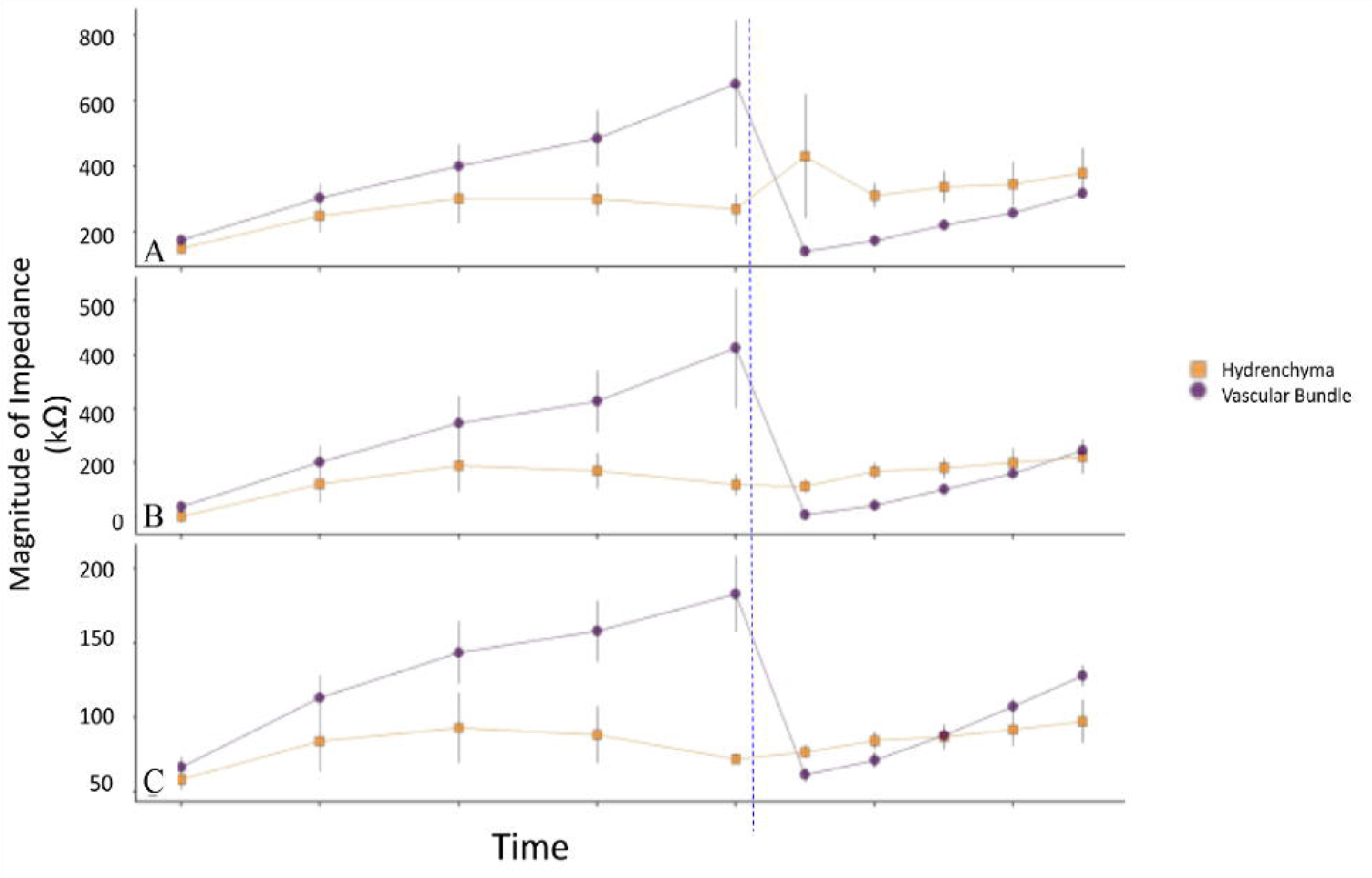
Magnitude of Impedance from 9am – 4pm. The mean impedance values of arrays on *S. bicolor* midribs. Magnitude of impedance of the abaxial and adaxial surface of a *S. bicolor* midrib from 9am to 3:30 pm at **(A)** 0.1 kHz; **(B)** 1 kHz; and **(C)** 10 kHz. Error bars represent 1 standard error, n=5. The orange represents EIS measurements collected in the hydrenchyma (H), and the purple in the vascular bundle (VB). Ticks on the x-axis represent 1 hour, starting at 9 am. The blue line represents the timing of irrigation.

**Figure 5:**
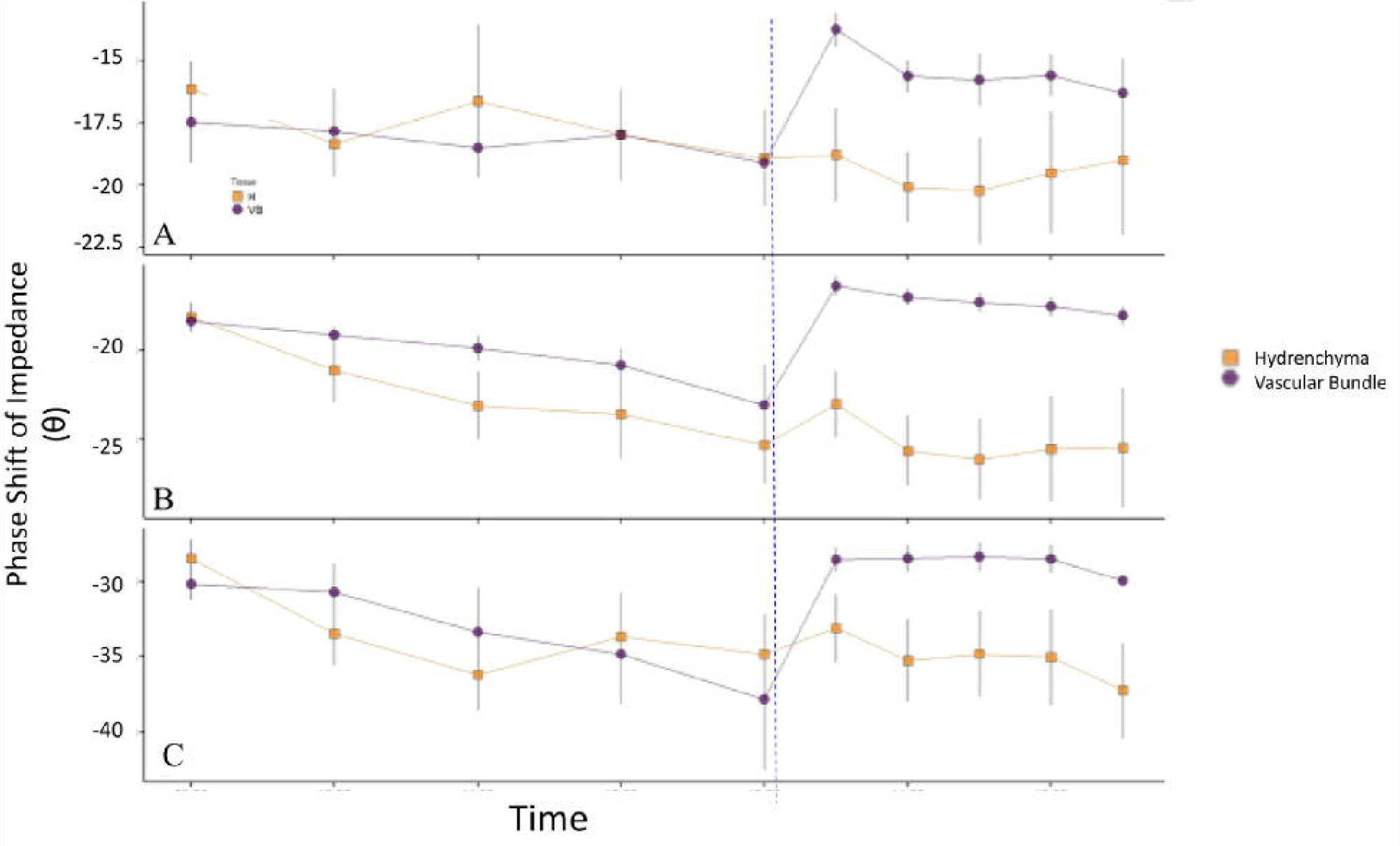
Phase Shift of Impedance from 9am – 4pm. Phase shift of impedance of the abaxial and adaxial surface of a *S. bicolor* midrib from 9am to 3:30 pm at **(A)** 0.1 kHz, **(B)** 1 kHz, and **(C)** 10 kHz. Error bars represent 1 standard error, n=5. The orange represents EIS measurements collected in the hydrenchyma (H), and the purple in the vascular bundle (VB). Ticks on the x-axis represent 1 hour, starting at 9 am. The blue line represents the timing of irrigation.

**Figure 6:**
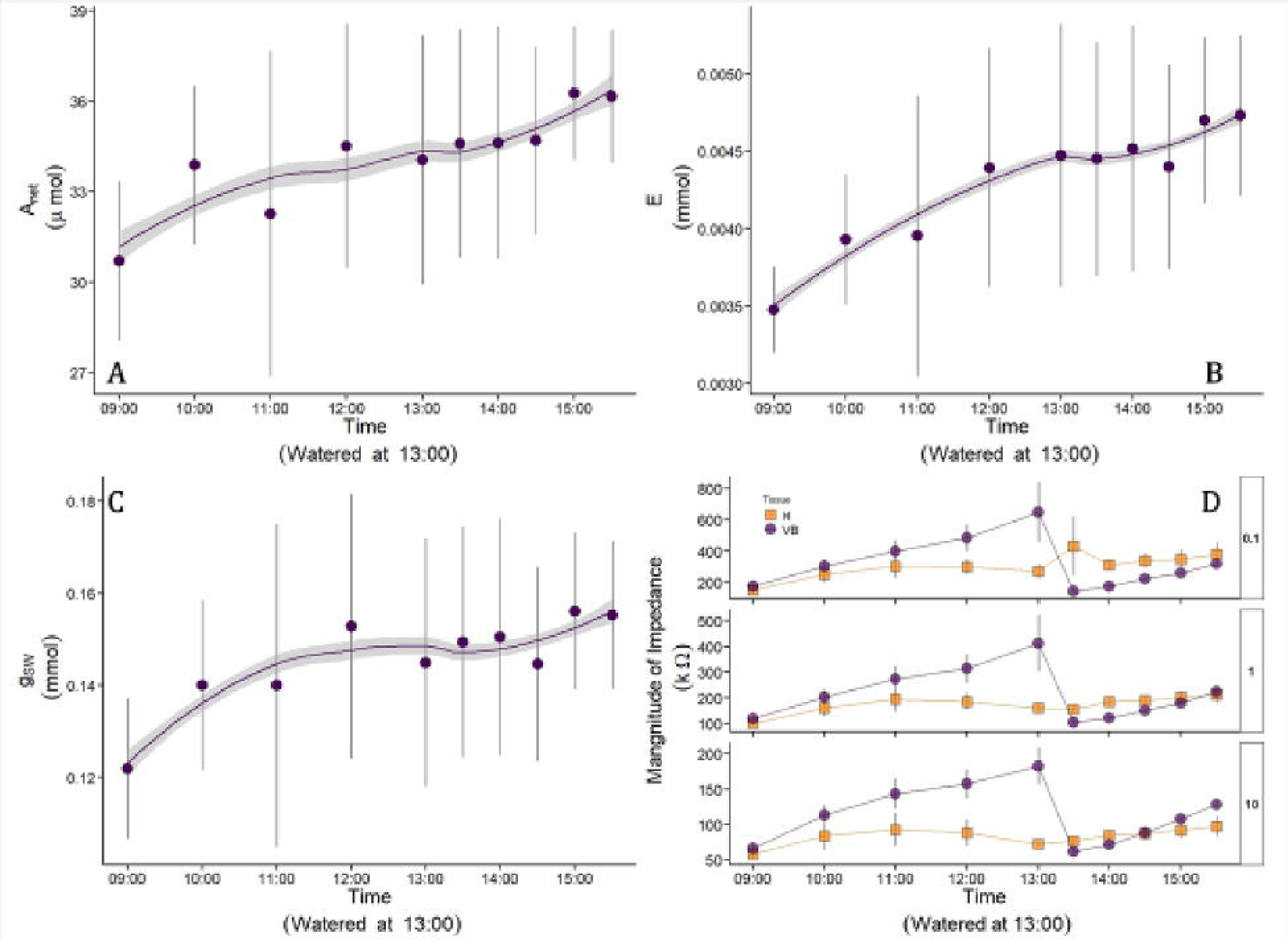
Gas Exchange with Magnitude of Impedance from 9am – 4pm. The mean gas exchange of *S. bicolor* leaves. Mean values of net assimilation (*A_net_*) in μmol; **(B)** transpiration (*E*) in mmol; **(C)** stomatal conductance (g*_sw_*) in mmol m^-2^s^-1^; **(D)** Magnitude of impedance (Orange – hydrenchyma (H); Purple – vascular bundle (VB)) in kOhms. Error bars represent 1 standard error, n=5. The collection time is on the x-axis.

Post water introduction, an immediate drop in abaxial impedance was recorded at each frequency, while the adaxial remained stable (p > 0.5 for all pairwise comparisons between 4hr 4.5hr in the hydrenchyma at all frequencies) (Figure 4). The reduction of impedance magnitude with the abaxial measurement returned to the 0hr magnitude immediately following water introduction. Planar transducer measurements continued to show random fluctuation in magnitude, which didn’t correlate with either morning water deprivation or mid-day watering (Supplemental Figure S4). Post water introduction, an immediate drop in abaxial impedance was recorded at each frequency, while the adaxial measurement remained stable (p > 0.5 for all pairwise comparisons between 4hr and 4.5hr in the hydrenchyma at all frequencies) (Figure 4). The reduction of impedance magnitude with the abaxial measurement returned to the 0hr magnitude immediately following water challenge. Planar transducer measurements continued to show random fluctuation in magnitude, which didn’t correlate with either morning water deprivation or mid-day watering.

Patterns seen in the impedance magnitude plots are reflected in the magnitude of impedance plots where the abaxial surface has consistent increases in resistance while the adaxial surface remains largely constant (Supplemental Figure S4). The increases in impedance on the abaxial measurement at lower frequencies were highly linear with R^2^ values of 0.99 at 0.1 kHz and 1 kHz between 900 hr and 1300 hr.

Phase angle (Θ) measurements were taken in tandem with impedance magnitude measurements at the insertion site-specific impedance responses (Figure 5). Phase angle data showed similar trends in initial phase angles at each frequency for abaxial impedance magnitude measurements which were responsive to the initial water deprivation and water challenge (Figures 3 & 5). Adaxial (hydrenchyma) phase angle measurements also were not responsive to the water introduction (Figure 5A).

### Gas Exchange Measurements

Gas exchange measurements were performed in tandem with the microneedle impedance measurements to determine if additional insight into the insertion site specific response could be gathered from orthogonal characterization (Figure 6). Assimilation (A), transpiration (E), and stomatal conductance (gsw) were measured on the same leaf as the microneedle impedance measurements as seen in Figure 4 (Figure 6A, B, C, & D respectively). The three gas exchange metrics showed patterns of increasing values until midday, then maintaining stability throughout the remainder of the experiment, however the error on the gas exchange measurements is greater than the change in values throughout the day.

## Discussion and Conclusions

In this study, we investigate if metal microneedles could act as microprobes for electrochemical impedance spectroscopy measurements of specific plant tissues. We elected to utilize simplistic approaches for adhering microneedles to the plant and for the choice of impedance analyzer as a proof of concept demonstration of field applicability. A representative illustration of the measurement methodology is seen in Figure 1A. Many previous EIS studies investigating plant impedance signals generated equivalent electrical circuit models and used those models for correlation of raw data into changes in their biologically representative circuit (Azzarello et al., 2012). In our approach, the hand-held LCR meter can only operate at 4 frequencies, which isn’t a sufficient number for creation of the frequency sweep (∼200) needed to generate an equivalent circuit model. Since the microneedle probes were hypothesized to selectively be positioned within a particular leaf anatomy, we continued under the assumption that this reduction of tissue components (i.e. epidermis, waxy cuticle) would simplify the tissue under investigation and enable high-resolution tissue interrogation in a field-like setting independent of a circuit model.

An initial puncture test of microneedles into sorghum midrib demonstrated the strength of the microneedles and they were regularly re-used throughout the course of this investigation. The microneedles even withstood washing and light polishing following continuous measurements to remove waxy plant leaf exudates. Overtime, some rounding of the microneedle tip was seen, but didn’t correlate to a shift in EIS performance. The rigidity of the midrib provided an unanticipated benefit for training new team members to apply the microneedle arrays. Breaching of the microneedles into the midrib produced a small, but well-defined, sensation. Since placement of the needles could be visually blind depending on the leaf orientation and desired surface, this sensation simplified placement of the device in the tissue of interest.

Initial impedance measurements comparing microneedle probes and planar probes confirmed data from other impedance reports in plants that removal of the epidermis greatly reduces the overall impedance magnitude of the measurement (Repo et al., 2000; Mancuso et al., 2004), although these studies used exclusively planar probes. While EIS has been used to correlate stem signals to environmental changes, to our knowledge, the utility of the in-situ approach for tracking water status changes within different tissues in any plant organ has not been investigated (Jeon et al., 2017; Garlando et al., 2022). Given the size and delicacy of a leaf, attaching the microneedle, with clip, to the base of the leaf midrib didn’t disrupt leaf function or result in significant damage to the leaf over the course of measurement (Figure 4).

While impedance spectroscopy provides a non-destructive methodology for studying biological systems, correlating impedance data to an underlying physiological cell or tissue change can be challenging (Azzarella et al., 2012). Orthogonal plant tissue scale monitoring methods do not exist thus further complicating data analysis. We can draw some conclusions regarding the in-situ measurement based on puncture tests and cross-sectional images in Figure 3. These experiments show that microneedle placement on one side of a leaf midrib does not reach needles placed on the opposing side of the leaf midrib. It is reasonable to assume that propagation of impedance signals occurs in the anatomical structures associated with the aspect of the leaf on which they were inserted. The distinguishing anatomical feature between the two surfaces is the presence of vascular bundles along the abaxial surface. Vascular bundles in mature leaves were measured to be ∼200 μm and are spaced ∼500 μm apart (Figure 1). As the microneedles used in this study were 220 μm wide at their base, it can be assumed that microneedle placement could be either between or within bundles. The consistent nature of abaxial impedance magnitude values at the start of the continuous monitoring experiments suggests impedance differences between those placements are negligible. This may be caused by the magnitude of impedance of the tissue under investigation. When current flows through an ideal conductor (<1 Ohm), it propagates via the least path of resistance and will not deviate much. Since the impedance magnitude of the midrib ranges from ∼50-200kOhm (at 0hr), depending on frequency, the impedance measurement may be averaging across a wider cross-sectional width of tissue. Additionally, the spacing (6cm) between EIS probes may also contribute to the wider field of signal propagation and ultimately, the tissue under measurement. Capitation or other physiological damage to the vascular bundle (including from the microneedle probes) would likely also change the signal propagation, such that the electrical signal is following the path with the least resistance; however, further investigation is needed to understand that dynamic.

Due to the challenges, interpretation of the time course data was performed by correlating insertion site-specific impedance data with the known anatomical location and function of tissues within the leaf midrib, and their assumed electrical properties through an irrigation experiment. Similar values of impedance and phase angle at the beginning of the time course experiment were interpreted as the two tissues being in equilibrium at a fully hydrated state. Visually, the plants appeared to be hydrated in that their leaves were full and not experiencing wilting. Cross-sectional images revealed the presence of hydrenchyma-like cells surrounding vascular bundles on the abaxial surface. If the microneedles positioned on the abaxial surface were not placed in a vascular bundle, they would be in these cells. However, over the course of measurement only the abaxial surface provided a temporal response to watering. Withholding water brought increases in the magnitude of impedance on the abaxial surface. This could be attributed to localized water stress due to the proximity of the sensors to vascular tissues, which would be the first tissues to experience decreased water flow from the soil.

All plants showed a similar pattern of impedance characteristics in the abaxial surface (vascular bundle) throughout the day, with a baseline measurement of about 400 kΩ at 0.1kHz. Impedance increased until the plant was watered, when it decreased and then subsequently started rising to meet pre-watering values (Figure 8a). The pattern was consistent across all measuring frequencies (Figure 8a, b, c). There was a significant impedance reduction between values at 1300 and 1330 (plants were watered after the 1300 measurements). Because the vascular bundles transport water from the roots to the leaves, this suggests the change in resistance values from 1300 to 1330 is an effect of the watering. Further studies will be required to determine the physiological response to watering that is causing the decrease in impedance. The changes may be caused by ionic solutes being transported through the vascular bundles, or cells surrounding the vascular bundle increasing intracellular transport.

Resistance values on the adaxial surface (hydrenchyma) increased by 12% from 900 to 1500 at 0.1kHz (Figure 8A, B, & C) and phase angle measurements did not appear to respond to watering at any of the frequencies (Figure 9 A, B, & C). The small change in the hydrenchyma may suggest that the plants were withdrawing water from the tissue, or that other signaling was occuring. Both the different values throughout the day and the different pattern, as well as the anatomical functions indicate the microneedle-based method is measuring two different tissues.

The vascular bundles also showed temporal variations in phase shift at 1 and 10 kHz, while the hydrenchyma showed little variation (Figures 8 & 9; B & C). At 0.1 kHz, both the hydrenchyma and vascular bundles showed variation with a clear pattern in response to watering (Figures 8 & 9; A). The phase shift data show that the vascular bundles became more capacitive throughout the day, until the plant was watered (Figure 9). After watering, the vascular bundles initially became more resistive, then returned to the pattern of increasing capacitance. Such a reduction in the magnitude in the low frequency measurements may provide better insight into extracellular impedance dynamics due to our understanding of frequency based cellular pathways (Figure 3b), which follow extracellular pathways at low frequency and transmembrane pathways at high frequency.

The insertion site-specific impedance response prompted additional investigation to determine the source of impedance fluctuation in the abaxial surface. Due to the initial one-hour equilibrium the plants experienced prior to measurement, thermal gradients across the thin leaf tissue (∼2mm thick) under measurement were assumed to be nearly constant. Puncture kinetics were similar for both sides of the leaf midrib, so tissue-needle interactions were also assumed to be consistent over the short time interval of this experiment. Further investigation into physiological changes and the measured electrical outputs present in xylem and phloem (embolism, p-plug damage, targeted herbivory of phloem, etc) would provide a more complete understanding of the water transport dynamics of the vascular bundles in field-grown crops.

Researchers have shown similar trends of increasing impedance during periods of drought, which is largely attributed to increasing values of the real resistance component of impedance, indicating that changes in the extracellular space are driving the fluctuations (Muramatsu and Hiraoka, 2007). A similar insertion site-specific impedance trend was seen after the 1300hr measurement with the abaxial measurement increasing in impedance while the adaxial surface remaining largely unchanged. Previous work studying heat stress in potato tubers showed near-constant values of resistance and capacitance until the temperature reached a threshold value (>40C) where a substantial decrease in extracellular resistance and membrane capacitance correlated with heat stress (Zhang et al., 1993). Due to the temperature control within the growth chamber, the primary increase in abaxial impedance does not appear to be attributed to heat stress and may be solely water dependent. Additionally, the standard errors of the planar arrays over the course of the day show no difference between the two surfaces and no response to watering. These results indicate the combined cuticle and epidermis act as electrical insulators, adding significant challenge to surface impedance measurements. This highlights the advantage of our microneedle technique for achieving subdermal electrode placement.

The lack of significant, measurable change in transpiration and photosynthesis after watering suggests the plant was not experiencing strong drought stress, and that the impedance data may detect pre-symptomatic water status. These results, in conjunction with changes in phase shift, suggest the decrease in magnitude of impedance in the vascular bundle does not simply represent transpiration, but rather, a systemic change in the tissue, either caused by microcavitation (small air pockets) in the xylem elements in response to the availability of water, activity of intracellular ion pumps in the surrounding cells, or permeability of transmembrane proteins (aquaporins). However, the measurement error on the gas exchange measurements was greater than the changes between data points (Figure 10).

Jeon et al. (2017) used microneedles as impedance probes in tomato stalks and a single needle contained both the impedance signal transmission and signal receiver on a planar microneedle. In this configuration, measurements of sap directly in contact with the transducers informed local salinity values for nutrient monitoring applications. Two distinct differences separate our microneedle-based approach from theirs. By placing the impedance signal transmission and signal receiver on two spatially separated microneedle arrays, the signal is forced to propagate through plant tissue instead of potentially only sap. Additionally, planar microneedles are in-plane with their substrate and can cause not perpendicular insertion angles, which in plants may create voids between the tissue and transducers that could cause loss of signal. Garlando et al. correlated water availability with EIS changes using needles that penetrated through the stem of a plant near the soil, but did not attempt to measure differences in tissues.

To our knowledge, this is the first study to demonstrate the ability to measure two tissues within a leaf, and that the two tissues will respond differently to subtle deviations in water status and water availability. Puncture tests and cross-sectional images (Figures 3 & 4) show that microneedle placement on one side of a leaf does not reach needles placed on the opposing side of the leaf. It is reasonable to assume that propagation of impedance signals occurs in the anatomical structures associated with the aspect of the leaf on which they were inserted. The distinguishing anatomical feature between the two surfaces is the presence of vascular bundles along the abaxial surface. Vascular bundles in mature leaves span about 200 μm, and are separated by ∼ 500 μm (Figure 1a). As the microneedles used in this study were 220 μm wide at their base, it can be assumed that microneedle placement could be either between or within bundles. The consistent nature of abaxial impedance values, which suggests impedance differences between those placements are negligible.

This study presents a novel application of a combination of microneedles and EIS as a wearable sensor that can monitor activity in two different tissues of the same leaf of a plant. Real-time monitoring of the disparate tissue responses to external factors, such as water availability, provides insight into the complex physiological functions of a leaf. The ability to monitor the impedance of the tissues without removing the leaf will allow for further studies of diurnal and seasonal changes. Establishing the ranges of EIS values in healthy plants may allow for more accurate detection of plant stress that has an adverse effect on the health and productivity of the plant.

## References

1. Azzarello, Elisa, Elisa Masi, and Stefano Mancuso. “Electrochemical impedance spectroscopy.” Plant electrophysiology: Methods and cell electrophysiology. Berlin, Heidelberg: Springer Berlin Heidelberg, 2012. 205–223.

2. Bragós, R., et al. “Biomass monitoring using impedance spectroscopy a.” Annals of the New York Academy of Sciences 873.1 (1999): 299–305.

3. Boyer, J. S. “Isopiestic technique: measurement of accurate leaf water potentials.” Science 154.3755 (1966): 1459–1460.

4. Boyer, JSf. “Leaf water potentials measured with a pressure chamber.” Plant Physiology 42.1 (1967): 133–137.

5. Ferreira, Javier, et al. “AD5933-based spectrometer for electrical bioimpedance applications.” Journal of Physics: Conference Series. Vol. 224. No. 1. IOP Publishing, 2010.

6. Greenwood, Duncan J., et al. “Opportunities for improving irrigation efficiency with quantitative models, soil water sensors and wireless technology.” The Journal of Agricultural Science 148.1 (2010): 1–16.

7. Jamaludin, Diyana, et al. “Impedance analysis of Labisia pumila plant water status.” Information Processing in Agriculture 2.3-4 (2015): 161–168.

8. Jeon, Eunyong, et al. “Development of electrical conductivity measurement technology for key plant physiological information using microneedle sensor.” Journal of Micromechanics and Microengineering 27.8 (2017): 085009.

9. Kim, Yeu-Chun, Jung-Hwan Park, and Mark R. Prausnitz. “Microneedles for drug and vaccine delivery.” Advanced drug delivery reviews 64.14 (2012): 1547–1568.

10. Koman, Volodymyr B., et al. “Persistent drought monitoring using a microfluidic-printed electro-mechanical sensor of stomata in planta.” Lab on a Chip 17.23 (2017): 4015–4024.

11. Levinsh, Gederts. “Water content of plant tissues: So simple that almost forgotten?.” Plants 12.6 (2023): 1238.

12. Mancuso, Stefano, et al. “Comparing fractal analysis, electrical impedance and electrolyte leakage for the assessment of cold tolerance in Callistemon and Grevillea spp.” The Journal of Horticultural Science and Biotechnology 79.4 (2004): 627–632.

13. Miller, Philip R., Roger J. Narayan, and Ronen Polsky. “Microneedle-based sensors for medical diagnosis.” Journal of Materials Chemistry B 4.8 (2016): 1379–1383.

14. Miller, Philip R., et al. “Fabrication of hollow metal microneedle arrays using a molding and electroplating method.” Mrs Advances 4.24 (2019): 1417–1426.

15. Muramatsu, Noboru, and Kiyoshi Hiraoka. “Water Status Detection of Satsuma Mandarin (Citrus unshiu Marc.) Trees using an Electrical Impedance Method.” Environmental Control in Biology 45.1 (2007): 1–7.

16. Repo, T., et al. “The electrical impedance spectroscopy of Scots pine (Pinus sylvestris L.) shoots in relation to cold acclimation.” Journal of Experimental Botany 51.353 (2000): 2095–2107.

17. Repo, Tapani, Janne Laukkanen, and Raimo Silvennoinen. “Measurement of the tree root growth using electrical impedance spectroscopy.” (2005).

18. Sishodia, Rajendra P., Ram L. Ray, and Sudhir K. Singh. “Applications of remote sensing in precision agriculture: A review.” Remote Sensing 12.19 (2020): 3136.

19. Wang, Ping M., et al. “Precise microinjection into skin using hollow microneedles.” Journal of investigative dermatology 126.5 (2006): 1080–1087.

20. van Iersel, Marc, et al. “Soil moisture sensor-based irrigation reduces water use and nutrient leaching in a commercial nursery.” Proc. Southern Nursery Assn. Res. Conf. Vol. 54. 2009.

21. Varlan, Anca Roxana, and Willy Sansen. “Nondestructive electrical impedance analysis in fruit: normal ripening and injuries characterization.” Electro-and Magnetobiology 15.3 (1996): 213–227.

22. Yin, Heyu, et al. “Soil sensors and plant wearables for smart and precision agriculture.” Advanced Materials 33.20 (2021): 2007764.

23. Zhang, M. I. N., et al. “Measurement of heat injury in plant tissue by using electrical impedance analysis.” Canadian journal of botany 71.12 (1993): 1605–1611.

24. Zhang, M. I. N., and J. H. M. Willison. “Electrical impedance analysis in plant tissues.” Journal of Experimental Botany 44.8 (1993): 1369–1375.

25. Zimmermann, D., et al. “A novel, non-invasive, online-monitoring, versatile and easy plant-based probe for measuring leaf water status.” Journal of experimental botany 59.11 (2008): 3157–3167.

## Software

26. R core Team (2018). R: A language and environment for statistical computing. R Foundation for Statistical Computing, Vienna, Austria. URL https://www.R-project.org/.

27. Hadley Wickham (2016). ggplot2: Elegant Graphics for Data Analysis. Springer-Verlag New York.

28. Hadley Wickham (2018). scales: Scale Functions for Visualization. R package version 1.0.0. https://CRAN.R-project.org/package=scales

29. Gert Stulp (2017). ggplotgui: Create GGplots via a Graphical User Interface. R package version 1.0.0. https://CRAN.R-project.org/package=ggplotgui

30. Simon Garnier (2018). viridis: Default Color Maps from ‘matplotlib’. R package version 0.5.1. https://CRAN.R-project.org/package=viridis

31. Garrett Grolemund, Hadley Wickham (2011). Dates and Times Made Easy with lubridate. Journal of Statistical Software, 40(3), 1–25. URL https://www.jstatsoft.org/v40/i03/

32. Stefan McKinnon Edwards (2018). lemon: Freshen Up your ‘ggplot2’ Plots. R package version 0.4.1. https://CRAN.R-project.org/package=lemon

33. Hadley Wickham (2011). The Split-Apply-Combine Strategy for Data Analysis. Journal of Statistical Software, 40(1), 1–29. URL https://www.jstatsoft.org/v40/i01/

34. Baptiste Auguie (2017). gridExtra: Miscellaneous Functions for “Grid” Graphics. R package version 2.3. https://CRAN.R-project.org/package=gridExtra

35. Douglas Bates, Martin Maechler, Ben Bolker, Steve Walker (2015). Fitting Linear Mixed-Effects Models Using lme4. Journal of Statistical Software, 67(1), 1–48. doi: 10.18637/jss.v067.i01.

36. Russell V. Length (2016). Least-Squares Means: The R Package lsmeans. Journal of Statistical Software, 69(1), 1–33. doi:10.18637/jss.v069.i01

37. Kuznetsova A, Brockhoff PB, Christensen RHB (2017). “lmerTest Package: Tests in Linear Mixed Effects Models.” Journal of Statistical Software, 82(13), 1–26. doi: 10.18637/jss.v082.i13 10.18637/jss.v082.i13

38. Torsten Hothorn, Frank Bretz and Peter Westfall (2008). Simultaneous Inference in General Parametric Models. Biometrical Journal 50(3), 346–363.

39. Ulrich Halekoh, Soren Hoisgaard (2014). A Kenward-Roger Approximation and Parametric Bootstrap Methods for Tests in Linear Mixed Models – The R Package pbkrtest. Journal of Statistical Software, 59(9), 1–30. URL http://www.jstatsoft.org/v59/i09/.

