## Supplemental Figures 1-4 for "Non-destructive monitoring of tissue-specific, irrigation responsive impedance signals in leaves using microneedles probes"

### Supplementary Figures

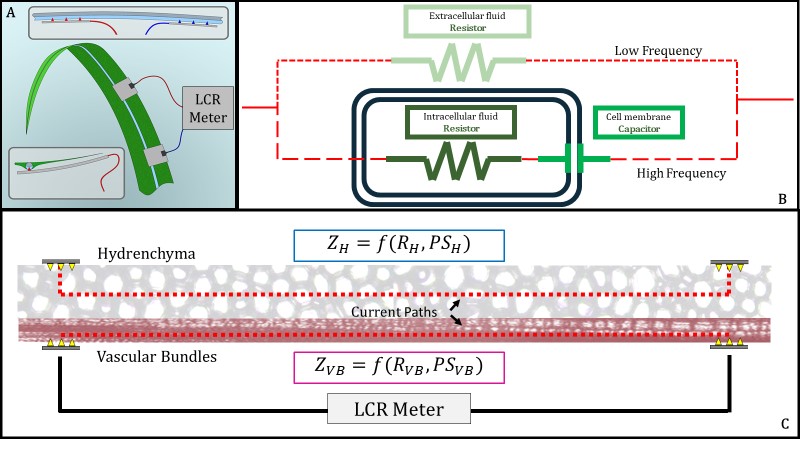

#### Supplemental Figure 1: Experimental design of the predicted electrical pathways

**(A)** Illustrated representation of tissue-specific impedance spectroscopy methods with microneedles in the midrib of a cereal crop connected to an LCR meter for measuring impedance characteristics of the tissue. The accompanying cross-section illustration demonstrates placement of two microneedles arrays along the midrib; **(B)** Hypothesized circuit diagram of a cell. In our proposed cell model, low frequency travels through the apoplastic pathway, through the extracellular fluid. High frequency travels through the cell membrane, through intracellular fluid. The cell membrane behaves like a capacitor. This circuit will be repeated in series throughout the tissue, between the two microneedle arrays; **(C)** Illustration of the proposed current pathways through the hydrenchyma and vascular bundles. By placing microneedles on the adaxial (top) surface of the midrib, current will travel through the hydrenchyma (H). Current can be directed through the vascular bundles (VB) by placing the patch on the abaxial (under) surface of the midrib. The LCR meter collects the resistance (R_x_) and phase shift (PS_x_), the respective real value and angle from the x-axis of impedance (Z).

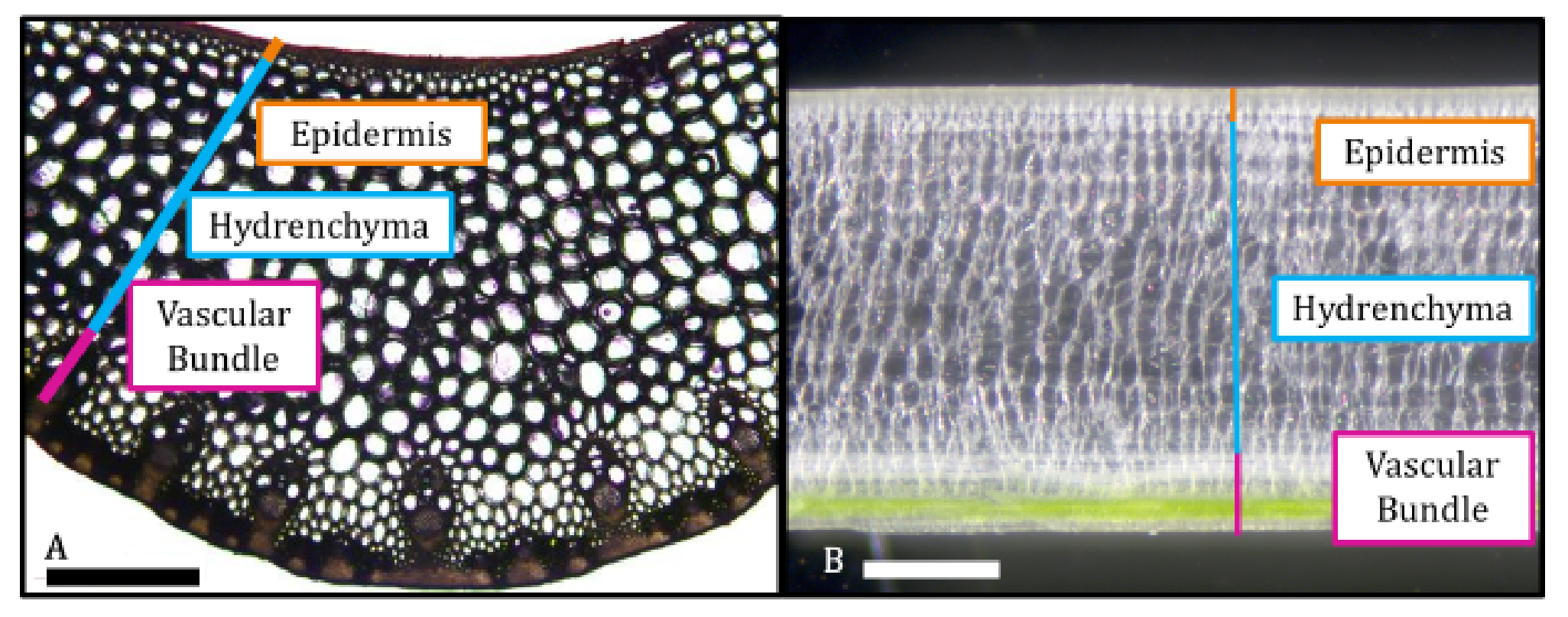

#### Supplemental Figure S2: *Sorghum bicolor* anatomy.

**(A)** Optical image of a *S. bicolor* sectioned perpendicular to the midrib show epidermal, hydrenchyma, and vascular tissue; **(B)** Optical image of a *S. bicolor* sectioned parallel the midrib show epidermal, hydrenchyma, and vascular tissue. Scale bars: 500um.

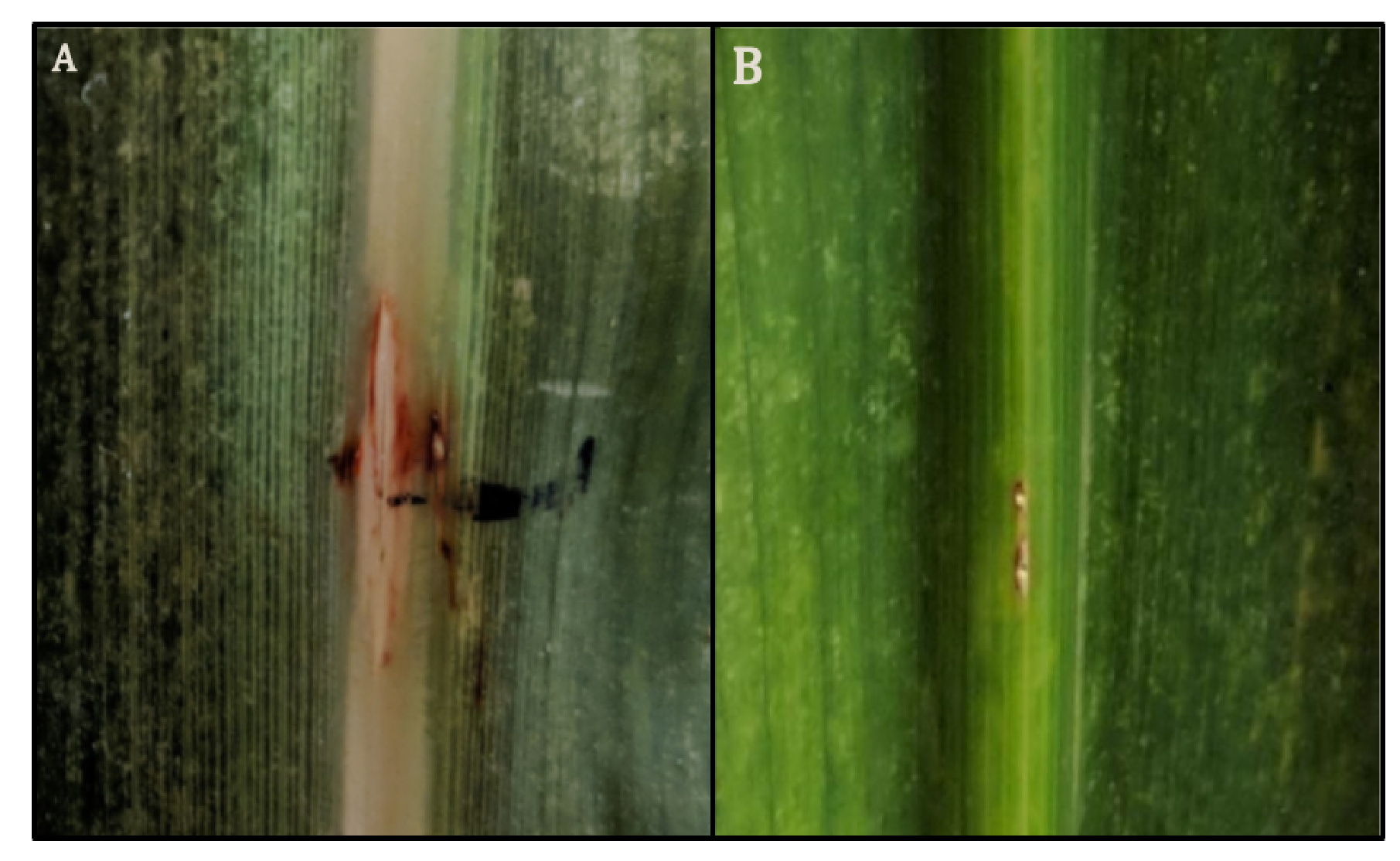

#### Supplemental Figure S3: Injection sites in the midrib for 14 days

**(A)** Adaxial (top) surface of the *S. bicolor* midrib. **(B)** Abaxial (bottom) surface of the *S. bicolor* midrib. These are representative images of 5 replicates. The adaxial image shows some splitting likely caused by a combination of the concave shape of the abaxial side of the mid-rib and the pressure of the clamps that were used to attach the needles.

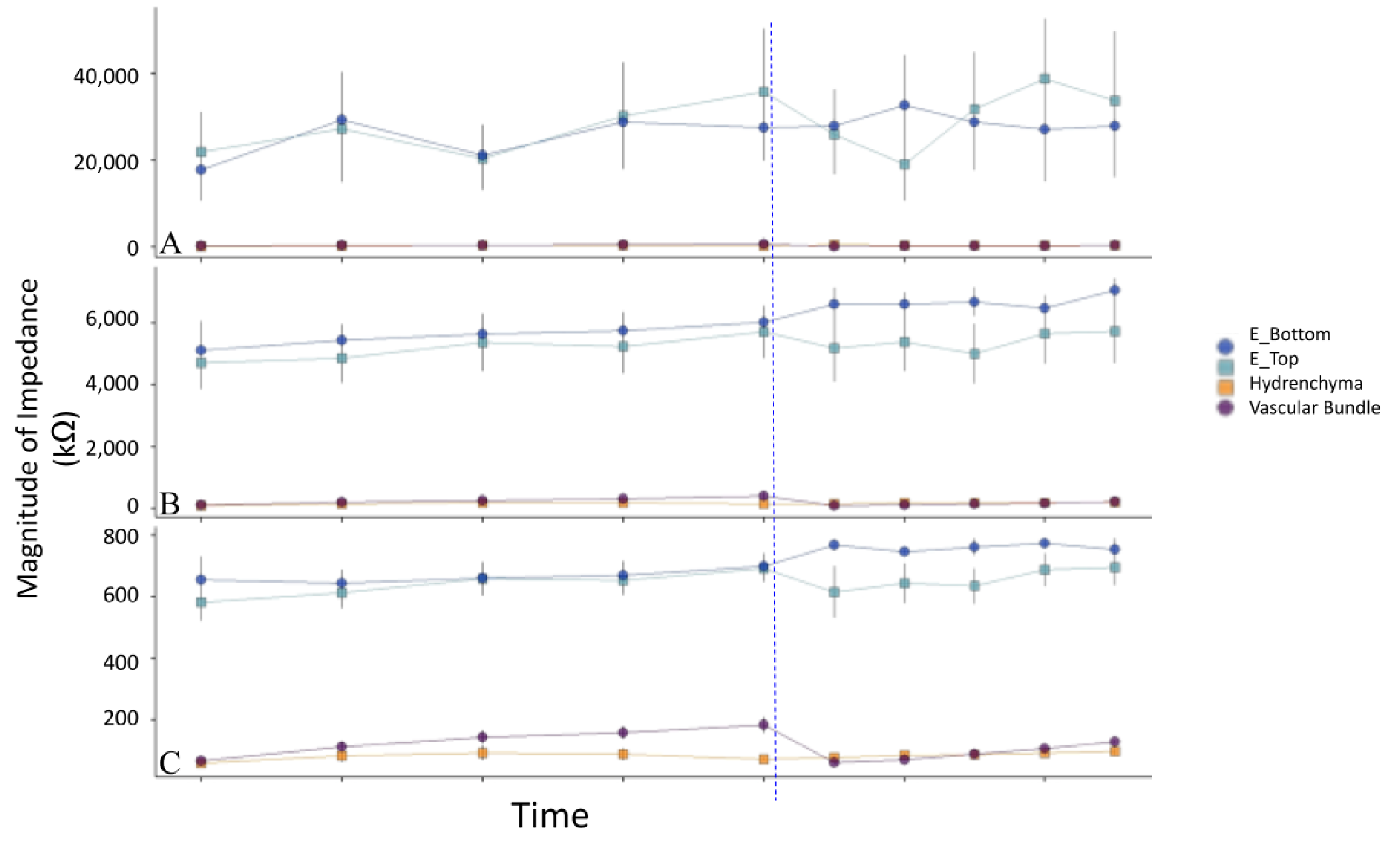

#### Supplemental Figure S4: Magnitude of Impedance with epidermal probes from 9am - 4pm

In kOhms (y-axis) throughout the measurement period (time on the x-axis). **(A)** magnitude of impedance at 0.1 kHz; **(B)** 1 kHz; and **(C)** 10 kHz. The blue represents EIS measurements collected through the epidermis (Dark blue - Bottom; Light Blue - Top). The orange represents EIS measurements collected in the hydrenchyma, and the purple in the vascular bundle. Error bars represent 1 standard error, n=5. Ticks on the x-axis represent 1 hour, starting at 9 am. The blue line represents the timing of irrigation.
